# Complex epistatic interactions between ELF3, PRR9, and PRR7 regulates the circadian clock and plant physiology

**DOI:** 10.1101/2023.07.17.547649

**Authors:** Li Yuan, Paula Avello, Zihao Zhu, Sarah C.L Lock, Kayla McCarthy, Ethan J. Redmond, Amanda M. Davis, Yang Song, Daphne Ezer, Jonathan W. Pitchford, Marcel Quint, Qiguang Xie, Xiaodong Xu, Seth J. Davis, James Ronald

## Abstract

Circadian clocks are endogenous timekeeping mechanisms that coordinate internal physiological responses with the external environment. EARLY FLOWERING3 (ELF3), PSEUDO RESPONSE REGULATOR (PRR9), and PRR7 are essential components of the plant circadian clock and facilitate entrainment of the clock to internal and external stimuli. Previous studies have highlighted a critical role for ELF3 in repressing the expression of *PRR9* and *PRR7*. However, the functional significance of activity in regulating circadian clock dynamics and plant development is unknown. To explore this regulatory dynamic further, we firstly employed mathematical modelling to simulate the effect of the *prr9/prr7* mutation on the *elf3* circadian phenotype. These simulations suggested that simultaneous mutations in *prr9/prr7* could rescue the *elf3* circadian arrythmia. Following these simulations, we generated all Arabidopsis *elf3/prr9/prr7* mutant combinations and investigated their circadian and developmental phenotypes. Although these assays could not replicate the results from the mathematical modelling, our results have revealed a complex epistatic relationship between ELF3 and PRR9/7 in regulating different aspects of plant development. ELF3 was essential for hypocotyl development under ambient and warm temperatures, while PRR9 was critical for root thermomorphogenesis. Finally, mutations in *prr9* and *prr7* rescued the photoperiod insensitive flowering phenotype of the *elf3* mutant. Together, our results highlight the importance of investigating the genetic relationship amongst plant circadian genes.

## Introduction

The daily rotation of Earth generates predictable *diel* cycles in light and temperature. Circadian clocks are molecular timekeeping mechanisms that anticipate these daily oscillations, allowing physiological responses to be coordinated with the external environment. This anticipatory ability is dependent on the circadian system undergoing daily re-setting in response to stimuli with predictable oscillatory patterns, a process called entrainment. In plants, light at dawn is thought to be a major entrainment signal (Millar, 2004). However, temperature, humidity and sugar availability also function in entrainment of the plant circadian clock (Webb et al., 2019). The circadian clock has a central role in the life history of plants, regulating germination, vegetative and floral development, metabolism, and the response to biotic and abiotic stress. As such, plants whose circadian clock is not closely aligned with the external environment have reduced fitness (Dodd et al., 2005, Greenham and McClung, 2015, Xu et al., 2022).

The plant circadian clock is an interconnected regulatory network of transcriptional, translational and post-translational feedback loops (McClung, 2019). Genetic and biochemical studies over recent decades have identified more than twenty different components that are involved in the circadian oscillator. Recent mathematical modelling has worked to reduce this complexity, resulting in a compact model that describes eight genes condensed into four components: CIRCADIAN CLOCK ASSOCIATED1 (CCA1) and LATE ELONGATED HYPOCOTYL (LHY) termed CL, PSEUDO RESPONSE REGULATOR9 (PRR9) and PRR7 termed P97, TIMING OF CAB EXPRESSION1 (TOC1) and PRR5 termed P51, and EARLY FLOWERING4 (ELF4) and LUX ARRYTHMO (LUX) termed EL (De Caluwé et al., 2016). Modifications of this simplified model have been used to understand the effect of external and internal cues on the circadian system and spatial differences in the plant circadian clock (Ohara et al., 2018, Avello et al., 2019, Greenwood et al., 2022).

In these compact models, the EL component describes the evening complex (EC). The EC is a tripartite protein complex composed of EARLY FLOWERING3 (ELF3), ELF4 and LUX (Nusinow et al., 2011, Herrero et al., 2012). LUX is a transcription factor that is necessary for the recruitment of ELF3 and ELF4 onto chromatin (Nusinow et al., 2011). ELF3 recruits chromatin remodeling enzymes including histone deacetylases, histone demethylases and nucleosome remodeling complexes to repress gene expression (Lee et al., 2019, Park et al., 2019, Tong et al., 2020, Lee and Seo, 2021). The role of ELF4 within the EC remains unresolved but may facilitate the nuclear localization of ELF3 and separately facilitate the binding of the EC to DNA (Kolmos et al., 2011, Herrero et al., 2012, Anwer et al., 2014, Silva et al., 2020, Ronald et al., 2021). Mutations in a single *ec* components result in circadian arrythmia, although the role and importance of the EC in regulating circadian rhythms in root cells is still uncertain (Covington et al., 2001, Doyle et al., 2002, Onai and Ishiura, 2005, Chen et al., 2020, Nimmo et al., 2020). In addition to regulatory activity in the oscillator, ELF3 is also necessary for light and temperature entrainment (Anwer et al., 2020, Zhu et al., 2022). ELF3 functions independently of the other EC components in mediating temperature entrainment (Zhu et al., 2022), while the requirement of the EC in facilitating ELF3-mediated light entrainment remains to be investigated.

A key target of the EC within the circadian clock are *PRR9* and *PRR7* (Kolmos et al., 2011, Herrero et al., 2012). PRR9 and PRR7 are transcription factors that share partial functional overlap within the circadian clock, with the *prr9*/*prr7* mutant having a longer circadian period than the respective single mutants (Farré et al., 2005, Salomé and McClung, 2005). Both PRR9 and PRR7 function in entrainment pathways of the circadian clock. The expression of *PRR9* is responsive to red light (RL) and this responsiveness is regulated by ELF3 (Farré et al., 2005, Ronald et al., 2022). *PRR7* expression is regulated by sugar availability and this regulation underpins sucrose entrainment of the oscillator (Frank et al., 2018). *PRR9* and *PRR7* also facilitate temperature entrainment, though the nature of this entrainment pathway remains to be investigated (Salomé and McClung, 2005). The EC directly regulates *PRR9* and *PRR7* expression (Kolmos et al., 2011, Herrero et al., 2012). At dusk, LUX binds to the promoter of both genes and ELF3 recruits the SWI2/SNF2-RELATED (SWRI) histone remodeling complex to induce a repressive chromatin state at both loci (Tong et al., 2020). Previous mathematical and genetic analysis indicates a stronger repressive effect of ELF3 on *PRR9* than *PRR7* (Herrero et al., 2012), although the significance of this remains unclear.

Although the expression of *PRR9* and *PRR7* is constitutively increased to a highly elevated level in the *elf3* mutant background (Kolmos et al., 2011), the importance of this mis-expression in contributing to the *elf3* circadian and physiological phenotypes has yet to be investigated. Here, through a use of mathematical modelling, molecular assays, and physiological measurements we sought to test the importance of the individual and high-order mutations in *prr9* and *prr7* in contributing to the different *elf3* phenotypes. Together, our results have highlighted a complex epistatic interaction between ELF3, PRR9, and PRR7.

## Results

### Modelling suggests a role of PRR9/PRR7 in the *elf3* arrythmia phenotype

To provide insights into the possible role of PRR9/PRR7 in contributing to the circadian phenotype of *elf3*, we firstly simulated the effects of the *prr9* and *prr7* mutations on the *elf3* circadian phenotype using the compact DC2016 model (De Caluwé et al., 2016). ELF3 is not implicitly modelled in the DC2016 model. Instead, the activity of ELF3 is represented by LUX and ELF4 within the EL component. *elf4* and *lux* mutants are similarly arrhythmic to *elf3* (Doyle et al., 2002, Onai and Ishiura, 2005). Hence, we will use simulated mutations in *el* as a proxy for mutations in *elf3*.

In the original compact model (DC2016), the *PRR9* (*P9*) and *PRR7* (*P7*) genes are grouped together in one component termed P97 (De Caluwé et al., 2016). Thus, it was firstly necessary to separate the P97 component into two components termed *P9* and *P7* (see materials and methods for further details). Three different models were then implemented; in the first model the individual P9 and P7 components retained the original functions of the P97 component as described in the DC2016 model. In the second model, we introduced a negative regulatory connection from CL (CCA1/LHY) to *P9*. In the third model, a negative regulatory connection from CL to *P9* and *P7* was introduced along with a negative auto-regulation in CL as described previously (Greenwood et al., 2022) (Figure 1A). The modifications were made to reflect CL now being described as repressors of *PRR* expression (Adams et al., 2015).

**Figure 1.**
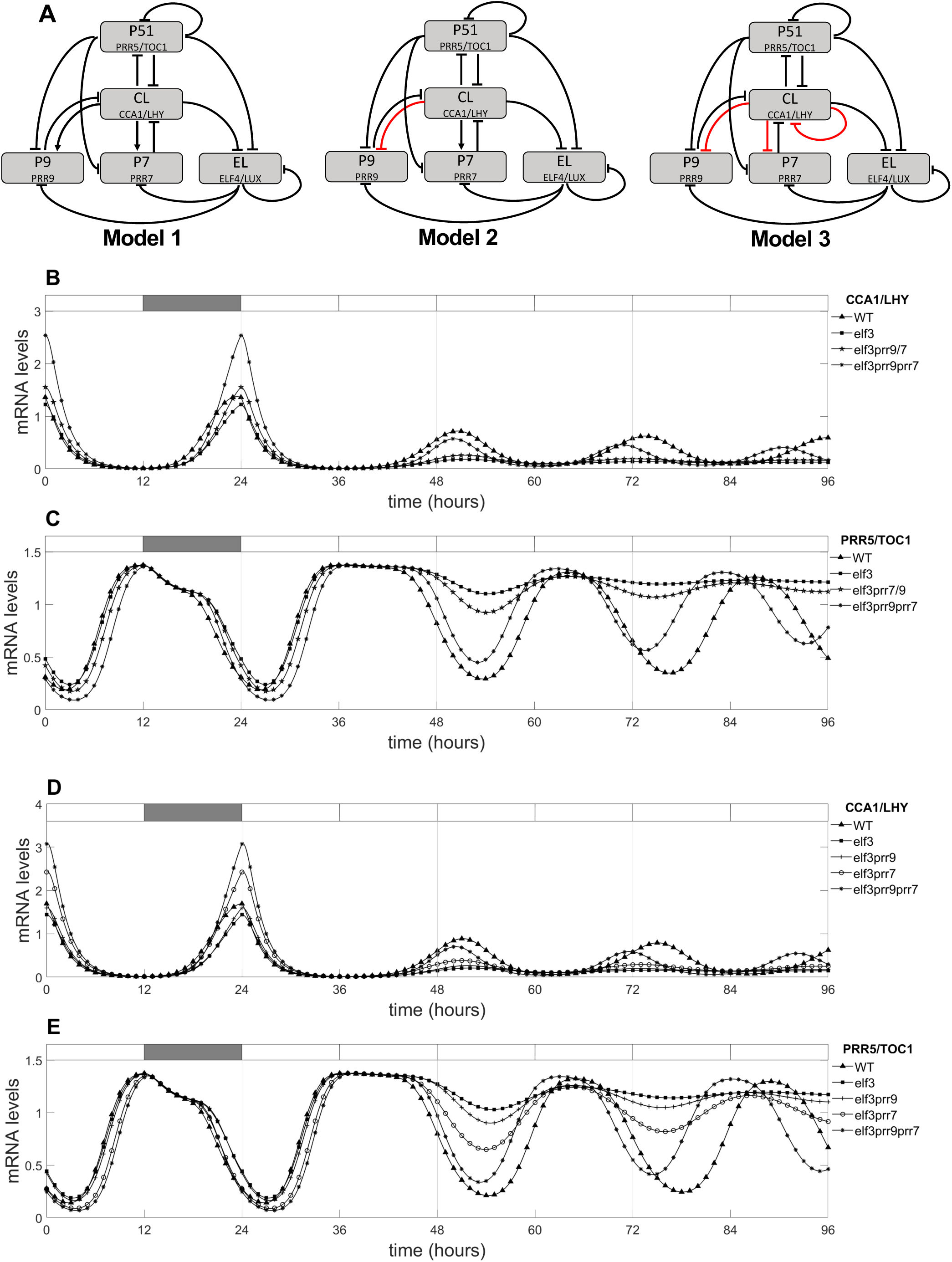
Modelling suggests that the *prr9/prr7* may contribute to the *evening complex* circadian arrythmia. (A) Three modifications of the De Caluwé *et al.,* 2016 (DC2016) model presenting *PRR9* and *PRR7* as separate components. Alterations to the original DC2016 model are highlighted in red. Model 1 is the original layout described in DC2016 but with P9 and P7 split into two components. In model 2, a negative interaction is introduced from CL to P9, while in model 3 a negative interaction is introduced from CL to P9 and P7, and negative auto-regulation of CL. Outputs from model 1 for the expression of (B) *CL*, and (C) *P51* and outputs from model 2 for the expression of (D) *CL* and (E) *P51* in the WT and simulated *elf3*/*prr* mutant backgrounds are shown. For model 1, the outputs for the *elf3*/*prr9* and *elf3/prr7* are represented as a single output (*elf3prr9/7*) as model 1 keeps the same functions for the respective mutants.

To understand whether the three different models could accurately simulate the *elf3/prr* phenotype we firstly simulated the expression of *CL* in wild type (WT) and mutants with well-defined circadian phenotypes. All three models were firstly conditioned for eight days of entrainment under neutral days (ND) photoperiods before being released into constant white light (CWL). The simulated expression of *CL* peaked at dawn for wildtype (WT) in model 1 and model 2, which is consistent with experimental data (Figure 1B, D) (Kolmos et al., 2011). The output of *CL* in the simulated *elf3*, *prr9*, *prr7*, and *prr9/prr7* mutants also closely replicated the reported expression profile of *CCA1/LHY* in the respective backgrounds for both model 1 and model 2 (Farré et al., 2005, Kolmos et al., 2011) (Figure 1B, D, Supplementary Figure 1A). In contrast, the expression of *CL* in model 3 did not replicate the reported expression profile of *CCA1/LHY* in WT or the different mutants (Supplementary Figure 2A). A similar behavior was also observed for *P51* (*PRR5/TOC1*). Again, the outputs of model 1 and model 2 closely followed the reported expression profile of *P51* for WT and the simulated mutants (Figure 1C, E, Supplementary Figure 1B) (Kolmos et al., 2011), while model 3 did not accurately reflect the expression of *P51* in any instance (Supplementary Figure 2B). Therefore, we will not further discuss the outputs of model 3 in this work.

We simulated the effect of combining the *elf3* and *prr9*, *prr7, prr9/prr7* mutations on the expression of *CL* and *P51* under free-running conditions. As with the *elf3* single mutant, the expression of *CL* was rhythmic in the *elf3/prr9* and *elf3/prr7* mutants under photocycles but became arrhythmic upon release into free-running conditions (Figure 1B, D). A similar behavior was observed for the output of *P51,* with the expression of *P51* quickly becoming arrhythmic upon release into free-running conditions for *elf3/prr9* and *elf3/prr7* (Figure 1C, E). In contrast, the expression of *CL* and *P51* remained rhythmic under free-running conditions in the simulated *elf3/prr9/prr7* mutant. Interestingly, the expression of *P51* and *CL* peaked earlier in the *elf3/prr9/prr7* than in WT in model 1 and model 2 (Figure 1B-D). Previous work has suggested that an extremely early phase in the *elf3* mutant may explain the *elf3* arrhythmic phenotype (Kim et al., 2005). Together, the output of these models suggests a role of PRR9 and PRR7 in the arrhythmicity of the *elf3* mutant.

### The p*rr9/prr7* mutations do not restore rhythmicity in the *elf3* mutant background

As the outputs of the models indicate PRR9/PRR7 may contribute to the arrhythmicity of the *elf3* mutant, we generated the different *elf3/prr9/prr7* mutant combinations to investigate this possibility. The *prr9-1/prr7-3* (*prr9/prr7* henceforth) double mutant was crossed into the *elf3-1 CCA1::LUC (elf3* henceforth) background to generate all *elf3/prr9/prr7* single, double and triple combinations with the *CCA1::LUC* reporter gene. This resulted in eight genotypic comparisons (Figure 2).

**Figure 2.**
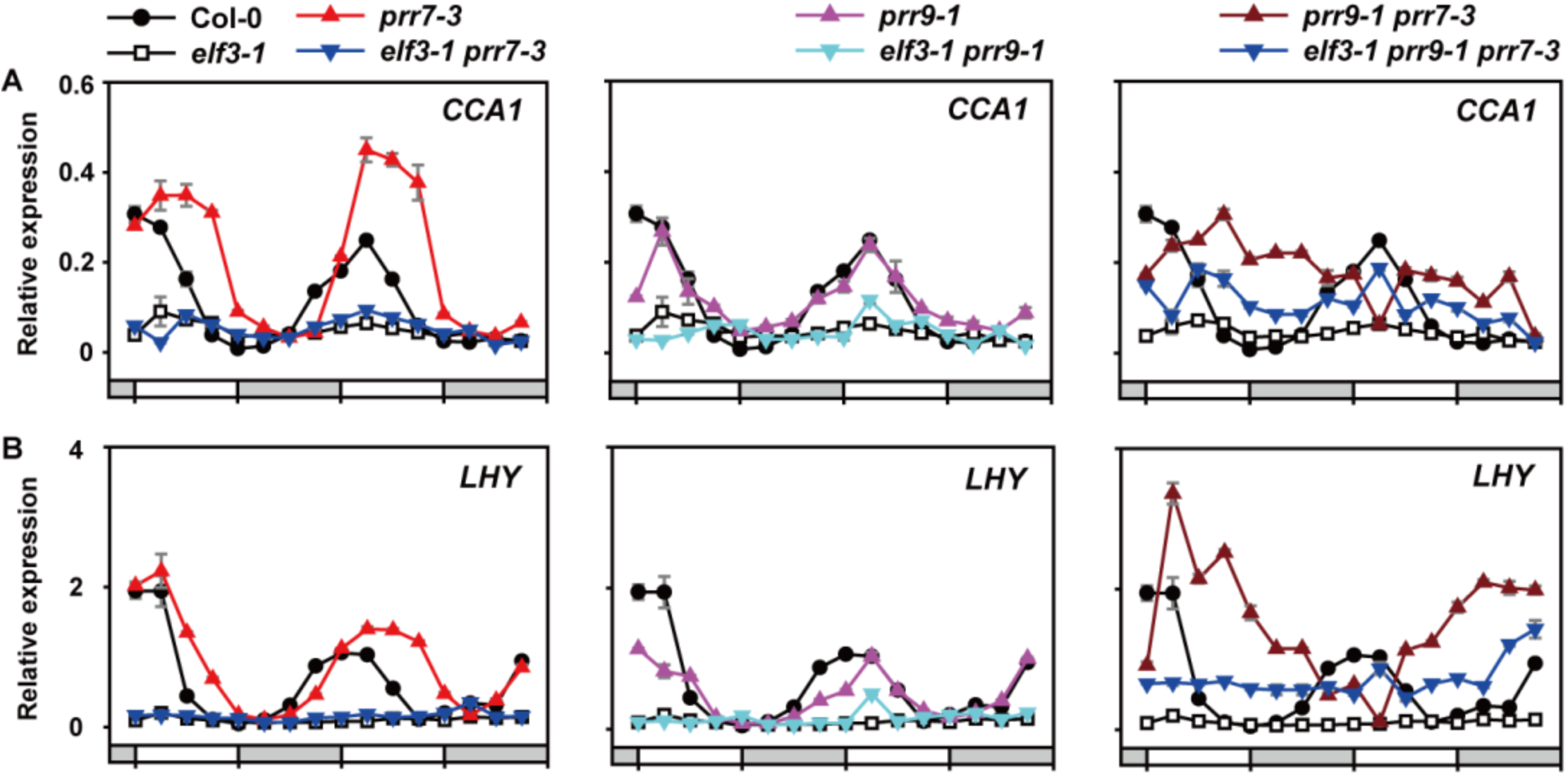
*prr9/prr7* does not rescue the arrhythmicity of *CCA1* or *LHY* expression in the *elf3* background. The expression of (A) *CCA1* or (B) *LHY* in wild type (Col-0), the *elf3*, *prr* and *elf3/prr* mutant backgrounds under constant light and constant temperature of 22°C. Seedlings were prior entrained under 12:12 light-dark cycles with a set temperature of 22°C for eight days. Data is the mean of three technical replicas, error bars represent standard error. All experiments were repeated twice, with similar results observed each time. *IPP2* was used as a normalization control. White or grey bars represent subjective day and subjective night.

We firstly investigated the expression of *CCA1* and *LHY* in the respective mutants entrained under a neutral day photoperiod and then released into constant light and constant temperature. As described previously, the expression of *CCA1* peaked at dawn in WT before rapidly declining (Figure 2A). In the *elf3* mutant, *CCA1* expression was arrhythmic and stayed suppressed below WT levels across the time course. There was no overt effect of the *prr9* mutation on *CCA1* expression, while in the *prr7* background the expression of *CCA1* was elevated and the peak accumulation was shifted from dawn to the early morning (Figure 2A). The peak expression of *CCA1* was further delayed in the *prr9/prr7* double mutant until the subjective afternoon. There was no noticeable effect of the *prr9* or *prr7* mutation on the *elf3* phenotype, with the respective double mutants closely resembling the *elf3* single and remaining arrhythmic (Figure 2A). The *elf3*/*prr9*/*prr7* triple mutant was also arrhythmic, but the expression of *CCA1* in the triple mutant was partially elevated relative to *elf3*. The expression pattern of *LHY* was comparable to that observed for *CCA1* for each respective mutant (Figure 2B). Neither the *prr9*, *prr7* or *prr9/prr7* mutations could rescue the arrhythmicity of *LHY* in the *elf3* background but, as before, there was a partial increase in the expression of *LHY* in the *elf3/prr9/prr7* triple mutant (Figure 2B). To confirm that neither the *prr9, prr7* or *prr9/prr7* mutant could rescue the *elf3* arrhythmic phenotype, we analyzed the output of the *CCA1::LUC* reporter. As with gene expression, all *elf3/prr* combinations were arrhythmic under free-running conditions (Supplementary Figure 3). In summary, our data suggests that the mis-expression of *PRR9* and *PRR7* cannot by themselves explain the arrhythmicity of the *elf3* mutant.

### ELF3 functions downstream of PRR9/7 in controlling ambient and warm-temperature induced hypocotyl elongation

Circadian regulation of hypocotyl elongation primarily occurs through the regulation of *PHYTOCHROME INTERACTING FACTOR4* (*PIF4*)/*PIF5* expression, stability, and transcriptional activity (Favero et al., 2021). Recently, ELF3 and PRR5/TOC1 were described to coordinately regulate the expression of *PIF4/5* in controlling hypocotyl elongation under SD and LD photoperiods (Li et al., 2020). Separate studies have also highlighted a role for PRR9/7 in regulating hypocotyl elongation via PIFs (Martín et al., 2018). Therefore, we tested whether there was a similar additive effect of the *elf3/prr9/prr7* mutation on hypocotyl elongation as described for the *elf3/toc1/prr5* mutant.

Firstly, we analyzed the hypocotyl phenotype of the *elf3/prr* mutants under short-day photoperiods with an ambient temperature of 20°C (Figure 3, Supplementary Figure 4). As reported previously, the *elf3* mutant had a long hypocotyl phenotype under these conditions compared to WT Col-0 (Figure 3A). There was no discernible hypocotyl phenotype in the *prr9* single mutant, while the *prr7* single mutant had a slightly longer hypocotyl compared to WT (Figure 3A). The hypocotyl of the *prr9*/*prr7* double mutant was further elongated than the *prr7* single mutant but remained shorter than the *elf3* single mutant. Notably, there was no effect of either *prr* single or double mutations on the *elf3* phenotype, with all mutant combinations closely resembling *elf3* (Figure 3A). Together, this suggests that PRR9/PRR7 redundantly regulate hypocotyl elongation and function in the same pathway as ELF3 rather than working cooperatively as previously described with ELF3 and TOC1/PRR5.

**Figure 3.**
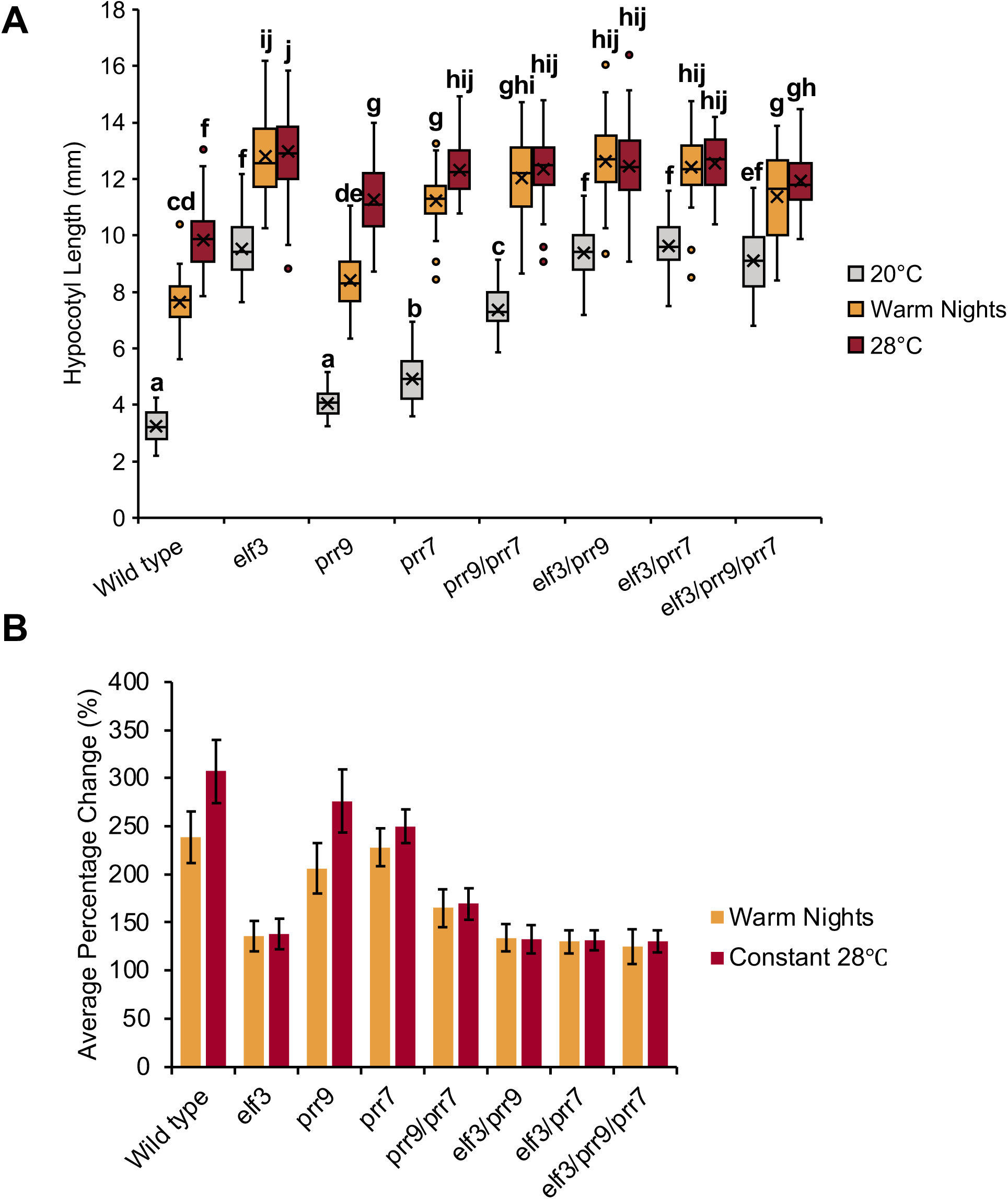
ELF3 functions downstream of PRR9/PRR7 in controlling hypocotyl development under ambient and elevated temperatures. (A) Hypocotyl length of wild type (Col-0) and the different *elf3*, *prr9* and *prr7* mutants under short-day photoperiods. Seedlings were grown with an ambient temperature of 20°C (grey), 28°C warm nights (orange) or constant 28°C (red) temperatures. (B) Average percentage change under warm nights or constant 28°C for the different genotypes. Error bars represent standard deviation. Experiments were repeated twice, with the presented data a combination of the two experiments. A minimum of 40 seedlings were analyzed. Letters signify a significant difference of p < 0.05, determined by ANOVA with a tukey HSD post-hoc test.

Next, we investigated the *elf3/prr* mutant hypocotyl phenotypes in SD photoperiods with either warm nights (28°C exclusively during the dark phase) or constitutive exposure to 28°C. In the WT Col-0 background, exposure to a 28°C warm-night was sufficient to promote hypocotyl elongation (Figure 3A). However, growth under constant temperatures of 28°C had a stronger effect on hypocotyl elongation compared to warm nights only (Figure 3B). The *prr9* and *prr7* single mutants had a similar response to WT, although there was a smaller difference between the constant 28°C and warm nights in the *prr7* background. Temperature responsiveness was strongly reduced in the *prr9/prr7* double mutant and there was no longer an additive effect of constant 28°C compared to warm nights (Figure 3B). Hypocotyl elongation in the *elf3* mutant was only weakly responsive to the elevated temperature, consistent with earlier reports (Jung et al., 2016, Ding et al., 2018). There was also no difference in response between warm nights or constitutive exposure to 28°C in the *elf3* background (Figure 3B). A similar response was also observed in the *elf3/prr* double and triple mutants, with all combinations closely resembling the response of the *elf3* single mutant (Figure 3A-B). Therefore, as with ambient temperature, *ELF3*, *PRR9* and *PRR7* likely function in the same pathway to control warm temperature induced hypocotyl elongation and *ELF3* functions downstream of *PRR9/PRR7* in this pathway.

### PRR9, but not ELF3, regulates regulate root development under warm temperatures

Unlike in the hypocotyl, the nature and role of the circadian clock in controlling root development continues to be unclear. Recent work highlighted a role for the EC in controlling lateral root development (Chen et al., 2020), while PRRs also regulates root development (Ruts et al., 2012, Li et al., 2019) and repress the expression of the *EC* components in root tissue under warm temperatures (Yuan et al., 2021). Therefore, we characterized the *elf3/prr* primary root phenotypes. As with hypocotyl development, we analyzed root growth under a SD photoperiod with either constant 20°C, 28°C warm nights only or constant 28°C. Root development was strongly impaired in the *elf3*, *prr9*, and *prr7* single mutants at 20°C compared to WT, with each respective single mutant having a similar response to each other (Figure 4A). There was no change in the primary root length of *prr9/prr7* double and *elf3/prr* double combinations compared to the respective single mutants. However, root development was further impaired in the *elf3/prr9*/*prr7* triple mutant compared to the single and double mutants (Figure 4A). Therefore, at ambient temperatures, ELF3 and PRR9/7 may regulate root development through separate but also partially overlapping pathways.

**Figure 4.**
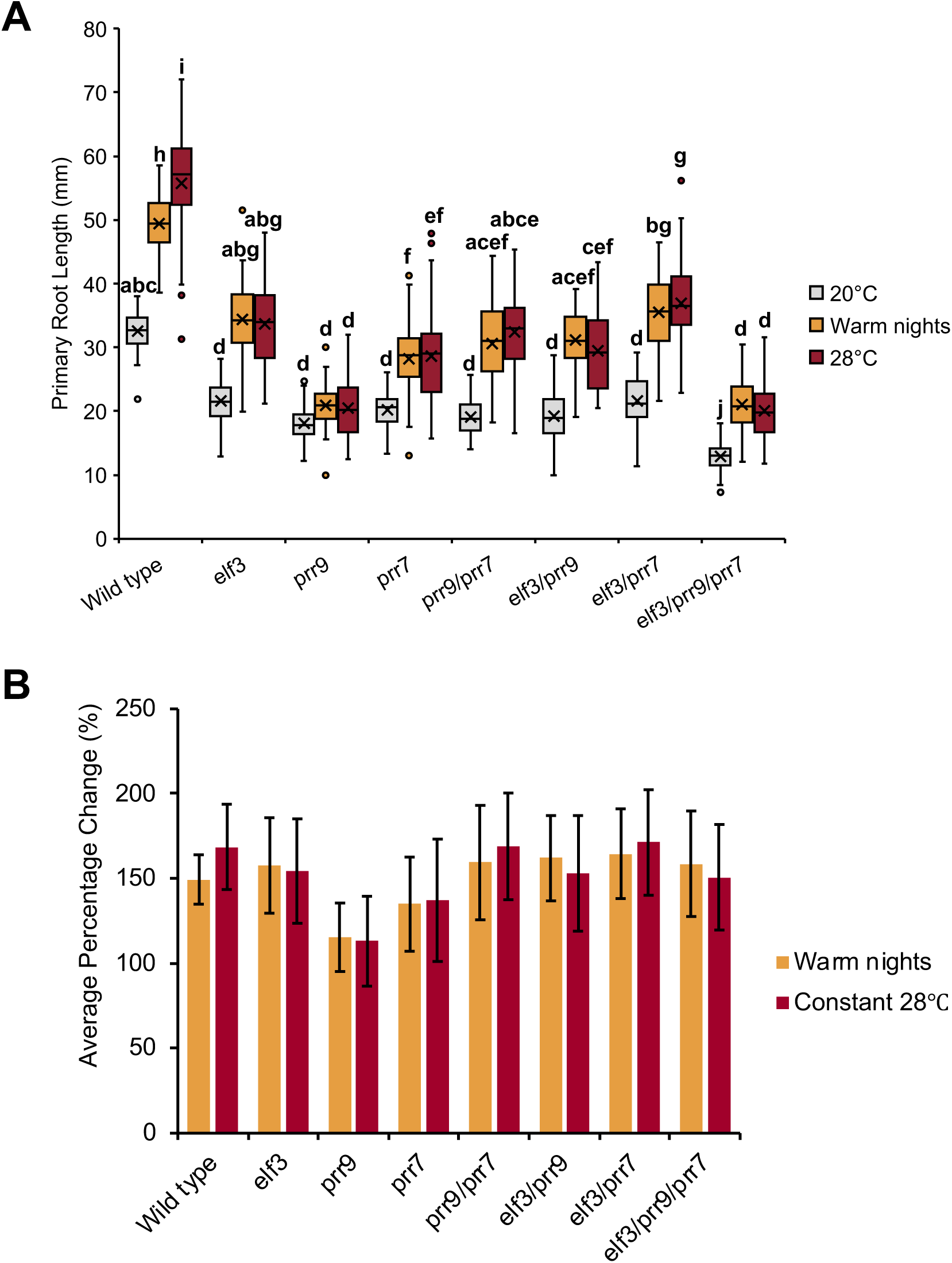
*PRR9* is necessary for root thermomorphogenesis. (A) Root length of wild type (Col-0) and the different *elf3*, *prr9* and *prr7* mutants under short-day photoperiods. Seedlings were grown with an ambient temperature of 20°C (grey), 28°C warm nights (orange) or constant 28°C (red) temperatures. (B) Average percentage change under warm nights or constant 28°C for the different genotypes. Error bars represent standard deviation. Experiments were repeated twice, with the presented data a combination of the two experiments. A minimum of 40 seedlings were analyzed. Letters signify a significant difference of p < 0.05, determined by ANOVA with a tukey HSD post-hoc test.

Exposure to warm temperature promoted root growth in WT, consistent with other reports in the literature (Quint et al., 2005, Hanzawa et al., 2013). As with hypocotyl development, constitutive exposure to 28°C caused a greater response than just warm nights, although the magnitude difference between the two growth conditions was smaller than observed for hypocotyl elongation (Figure 3B, 4B). Root growth in the *elf3* single mutant remained responsive to warmth (Figure 4A) but there was no difference in the magnitude of response between constant exposure to 28°C or warm nights only (Figure 4B). Root development was also thermoresponsive in the *prr7* mutant, but this response was weaker than the response seen in WT or the *elf3* mutant (Figure 4A-B). As with the *elf3* mutant, root development in the *prr7* mutant also did not show a differing response between the constant 28°C or warm night only (Figure 4B). In the *prr9* mutant, the temperature responsiveness of root growth was largely lost, with only a minimal, non-significant, response observed under both warm conditions (Figure 4A-B). Notably, thermo-responsiveness was restored in the *prr9*/*prr7* double mutant, suggesting a complex regulatory pathway underpinning the *prr9* and *prr7* phenotype. Root growth in the *elf3/prr* double and triple mutants was also thermoresponsive and showed a similar response to WT and the *elf3* single mutant (Figure 4A-B). However, as with the *elf3* mutant, there was no longer a difference in response between warm nights and constitutive exposure to 28°C (Figure 4B). Altogether, our results suggest a complex relationship between ELF3 and PRR9/7 in controlling root development in response to warm temperatures.

### *elf3* photoperiod insensitivity is rescued by the *prr9/7* mutation

ELF3 and PRRs are both critical regulators of the photoperiodic flowering time pathway in Arabidopsis and other plant species (Osnato et al., 2022). In Arabidopsis, mutations in *elf3* and *prr9/7* lead to opposite effects on flowering time; *elf3* mutants are early flowering and photoperiod-insensitive (Zagotta et al., 1996), while the *prr9/7* retains photoperiod sensitivity, but flowers late under LD and SD (Nakamichi et al., 2007). This differing response between the *elf3* and *prr9/7* mutant led us to explore potential epistatic interactions in flowering time under LD and SD.

Under LD, the *elf3* mutant flowered slightly earlier than WT plants, while the *prr9* and *prr7* single mutants had a moderate late-flowering phenotype (Figure 5A-B). This late-flowering phenotype was enhanced in the *prr9/prr7* double mutant, which flowered much later than respective single mutant (Figure 5A-B). Introducing the *prr9* mutation into the *elf3* background resulted in a similar response as to the *elf3* single mutant. For the *elf3/prr7* mutant, flowering time trended towards a WT response for both measurements of flowering (Figure 5A-B). However, there was no significant difference between *elf3* and *elf3/prr7* in the number of leaves produced. In the *elf3/prr9/prr7* background, flowering was delayed relative to WT as measured by days to flower but not leaf count (Figure 5A-B). Together, these results indicate that PRR9/7 and ELF3 likely regulate flowering time under LD through partially overlapping pathways, with PRR9/7 functioning downstream of ELF3.

**Figure 5.**
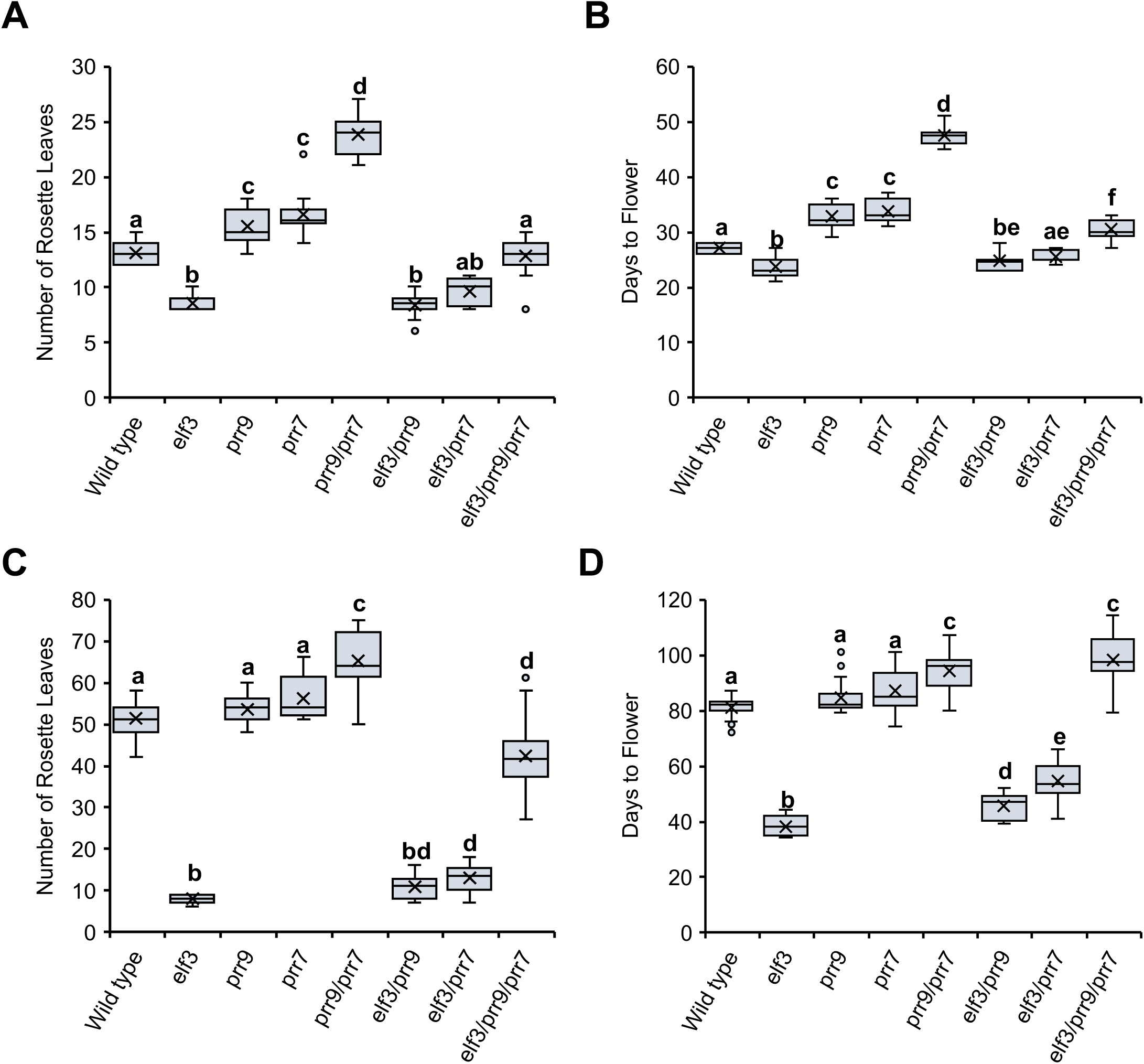
Mutations in *prr9/prr7* delays the *elf3* flowering phenotype under long and short-day photoperiods. Flowering time in the wild type (Col-0), *elf3*, *prr* and *elf3/prr* mutants under long-days (A-B) or short-days (C-D). Flowering was measured under each photoperiod by the number of rosette leaves (A, C) or the days taken to produce a bolt that was ∼1 cm above the rosette (B, D). For both photoperiods plants were grown under a constant temperature of 22°C. Experiments were repeated twice, with a minimum of eight plants analyzed for each experimental repeat. Letters signify a significant difference of p < 0.05 as determined by ANOVA with a tukey HSD post-hoc test.

Under SD, *elf3* mutants flowered extremely early (Figure 5C-D). Neither the *prr9* nor the *prr7* mutant had a flowering-time phenotype under SD, with both mutants flowering at the same time as WT. Flowering was delayed in the *prr9/prr7* double mutant, suggesting genetic redundancy between PRR9/7 in regulating flowering under SD photoperiods (Figure 5C-D). Both the *prr9* and *prr7* mutations were able to partially rescue the *elf3* early flowering phenotype under SD and, as with LD, the *prr7* mutation had a stronger effect on rescuing the *elf3* phenotype than *prr9* (Figure 5). Neither the *prr9* nor *prr7* mutation alone was able to fully restore the *elf3* phenotype to a WT response, with both *elf3/prr9* and *elf3/prr7* still flowering earlier than WT plants. The phenotype for *elf3/prr9/prr7* mutant was complex, with a disconnect between leaf number and days to flower (Figure 5C-D). As measured by days to flower, the *elf3/prr9/prr7* triple mutant resembled the *prr9/prr7* double mutant. However, when the number of leaves were measured, the *elf3/prr9/prr7* had fewer leaves than WT. Therefore, this analysis suggests a wider metabolic or plastochron phenotype in the *elf3*/*prr9*/*prr7* background.

## Discussion

PRR9 and PRR7 are direct targets of ELF3 through the activity of the EC (Herrero et al., 2012, Tong et al., 2020) and as a result the expression of *PRR9* and *PRR7* is constitutively upregulated in the *elf3* mutant background (Kolmos et al., 2011). However, the consequence of this mis-expression has so far not been tested. Here, we utilized mathematical modelling to simulate the effect of the *prr9* and *prr7* mutations on the circadian phenotype of the *elf3* mutant. The outputs of the modified DC2016 models indicated a possible role for *PRR9* and *PRR7* in contributing to *elf3* arrhythmicity (Figure 1). However, follow-up experiments analyzing the circadian phenotypes of the different *elf3/prr* mutant through gene expression analysis (Figure 2), or luminescence reporter (Supplementary Figure 3) revealed that all combinations of the *elf3/prr9/prr7* mutants were arrhythmic. It is unclear why the mathematical models and our *in planta* data produced different outputs. PRR9 and PRR7 are transcriptional repressors that are part of a gene family that includes *TOC1*, *PRR3*, and *PRR5.* So far, the EC has been demonstrated to directly repress the expression of *TOC1,* as well as *PRR9* and *PRR7* (Herrero et al., 2012, Lee et al., 2019). Whether the EC represses *PRR3* or *PRR5* expression remains to be investigated. It is possible that the mis-expression of additional *PRRs* in the *elf3* background also contributes to the circadian arrythmia phenotype and further *prr* mutations are necessary alongside the *prr9/prr7* mutations. However, equally, limitations within the implemented models may be responsible for the discrepancies we have observed, and it would be relevant to focus further modelling efforts on the understanding of these discrepancies for future investigation.

Alongside investigating the effect of the *prr9/prr7* mutation on the *elf3* circadian phenotype, we also characterized the *elf3/prr* developmental phenotypes. We firstly investigated hypocotyl development, where we found that the *elf3/prr* combinations closely resembled the *elf3* phenotype and there was no enhancement of the *elf3* phenotype (Figure 3). This suggests that PRR9/7 function upstream of ELF3 in the same genetic pathway to control hypocotyl development. Circadian regulation of hypocotyl development primarily occurs through the control of PHYTOCHROME INTERACTING FACTOR (PIF) activity. ELF3 regulates the expression of *PIF4*, *PIF5*, and *PIF7* through the EC, while independently also controlling PIF4 transcriptional activity (Nusinow et al., 2011, Nieto et al., 2015, Jiang et al., 2019). Separately, PRR9/7 regulate *PIF4* expression, while also directly inhibiting PIF4 functional activity (Liu et al., 2016, Martín et al., 2018). Thus, it was unsurprising that there is no further enhancement of the *elf3* phenotype in the *elf3/prr9/prr7* background as *PIF4* expression and activity is already enhanced in the *elf3* background. We also observed that the *elf3* and *prr* single mutants were equally compromised in root development (Figure 4). This short-root phenotype was not enhanced in the *prr9/prr7* or *elf3/prr* double mutants, with all lines displaying similar phenotypic defects as the respective single mutants. However, the *elf3/prr9/prr7* triple mutant had a shorter root than the respective single mutants, indicating that ELF3 and PRR9/7 independently control root development (Figure 4). So far, few studies have investigated how the circadian clock controls root development. PRR9/7 may regulate root development via controlling TOR signaling (Li et al., 2019), but further studies are needed to understand how the circadian clock controls root development.

Exposure to constitutively elevated temperature promotes hypocotyl and root development (Quint et al., 2016, Lee et al., 2021). Here, we observed that elevated temperatures during the night was also sufficient in promoting hypocotyl and root development in WT plants. However, hypocotyl and root development more strongly responded to constant exposure to warm temperatures than night-time only exposure to warm temperature (Figure 3, 4). ELF3 was necessary for elongation of the hypocotyl in response to warm temperatures regardless of the duration of temperature exposure (Figure 3B), supporting previous work (Box et al., 2015, Raschke et al., 2015, Zhang et al., 2021). In contrast, root development remained sensitive to temperature in the *elf3* mutant background but there was no difference between different durations of warmth exposure (Figure 4B). Together, our results support recent work that highlighted shoot thermosensors may not function as root thermosensors, and thermomorphogenesis in shoots and roots uses different genetic pathways (Lee et al., 2021, Borniego et al., 2022, Ai et al., 2023).

Hypocotyl elongation in *prr9* and *prr7* remained temperature responsive under both constant conditions and warm nights only, although there was a smaller magnitude difference between the two different conditions in the *prr7* mutant (Figure 3B). The temperature responsiveness of the *prr9/prr7* double mutant was further impaired and there was no magnitude difference between the different warm temperature regimes, indicating a redundant role for PRR9/PRR7 in mediating daytime hypocotyl thermomorphogenesis (Figure 3B). TOC1 regulates the time-of-day response to warm temperature by directly binding to and subsequently inhibiting the activity of PIF4 (Zhu et al., 2016). This mechanism ensures thermomorphogenesis is phased to occur only in the late evening and early morning. Under ambient temperatures, PRR9 and PRR7 directly interact with PIF4 to control the timing of *CYCLING DOF FACTOR6* (CDF6) expression, a positive regulator of hypocotyl elongation (Martín et al., 2018). Whether similar mechanisms control hypocotyl thermomorphogenesis remains to be investigated.

In the roots, our results suggest a complex genetic pathway underpins the response to warm temperature response. Root development in the *prr9* mutant was insensitive to warm temperature regardless of the length of warmth exposure (Figure 4B). This response was then fully rescued in the *prr9/prr7*, *elf3/prr9* and *elf3/prr9/prr7* mutant combinations. So far, no studies have highlighted a role for PRR9 in controlling root thermomorphogenesis and the pathway(s) regulating root thermomorphogenesis remain unclear. A recent study has revealed that root thermomorphogenesis is mediated by an unidentified regulator that controls auxin-dependent progression of the cell cycle (Ai et al., 2023). This signaling pathway occurs independently of the well described PIF thermal integration pathway that regulates thermomorphogenesis in aerial tissue (Ai et al., 2023, Delker et al., 2022). Thus, it is unclear how the circadian clock regulates root thermomorphogenesis. TOC1, another PRR within the circadian clock, has been described to regulate cell cycle progression in leaf tissue (Fung-Uceda et al., 2018). Alternatively, all stages of auxin homeostasis and signaling are under circadian control (Covington and Harmer, 2007). Further studies are needed to investigate the integration point for the circadian clock in controlling root thermomorphogenesis.

The regulatory connection between ELF3 and PRRs has emerged as critical node in the photoperiodic control of flowering time in agronomically important crops (Faure et al., 2012, Zhao et al., 2012, Alvarez et al., 2022, Woods et al., 2022). In Arabidopsis, the importance of the *PRRs* in contributing to the photoperiod-insensitive early flowering of the *elf3* mutant has remained untested. Under LD, introducing the *prr9/prr7* mutation into the *elf3* background rescued the early flowering *elf3* phenotype and resulted in a mild late-flowering phenotype when measured by days to flower (Figure 5B) but only a WT response when measured by leaf count (Figure 5A). These discrepancies were further enhanced under SD. When measured by days to flower, the *elf3/prr9/prr7* triple mutant resembled the late flowering *prr9/prr7* double mutant (Figure 5D). However, by leaf count the *elf3/prr9/prr7* triple mutant had fewer leaves than WT (Figure 5C), thus technically an early flowering plant. Together, these results may highlight a plastochron phenotype in the *elf3/prr9/prr7* mutant. Plastochron regulation is a complex trait, with multiple signaling pathways contributing to the timing of leaf emergence (Werner et al., 2001, Wang et al., 2008, Lohmann et al., 2010, Mimura et al., 2012, Cornet et al., 2021). One regulator of the plastochron is gibberellin (GA) signaling (Mimura et al., 2012, Mimura and Itoh, 2014). In barley, *elf3* mutants over-accumulate GA and this mis-regulation contributes to the *elf3* early flowering under non-inductive SD photoperiods (Boden et al., 2014). It will be of interest to investigate whether GA is a contributory factor to the complex flowering time phenotypes observed in the *elf3/prr9/prr7* mutant background.

### Concluding Remarks

The *elf3* circadian arrhythmicity was first described nearly 30 years ago (Hicks et al., 1996) but we still do not understand the causative factors. In this study, we hypothesized that mis-regulation of *PRR9* and *PRR7* may explain the *elf3* arrhythmicity. Although our mathematical modelling supports such a hypothesis (Figure 1), we could not replicate such results *in vivo* (Figure 2, Supplementary Figure 3). Further work is needed to untangle whether further genetic redundancy amongst the *PRRs* is responsible for the discrepancy between the simulations and *in vivo* results.

## Materials and Methods

### Plant Lines

All mutant lines described in this work are in the Col-0 background. The Col-0, *prr9-1*, *prr7-3* and *prr9-1/prr7-3* mutant lines harboring *CCA1::LUC* have all been described previously (Farré et al., 2005). The new mutant combinations generated in this work were created by crossing the *elf3-1* (Zagotta et al., 1996) *CCA1::LUC* mutant into the *prr9-1/prr7-3* double mutant. The various single, double, and triple mutant combinations were then identified through genotyping.

### Modelling

The original De Caluwé model considered *PRR9* and *PRR7* having similar expression profiles and they were characterized as a single model variable (P97) (De Caluwé et al., 2016). Here we modified the DC Caluwé model to separate the variable P97 so that three models were implemented:

▪ Model 1: The role of each separated component within the network, as described by De Caluwé et al. 2016, was unchanged.
▪ Model 2: The interaction between *CCA1/LHY* (CL) *and PRR9* (P9) was modified; *PRR9* is inhibited by *CCA1/LHY*.
▪ Model 3: The interaction between *CCA1/LHY* (CL) and *PRR9*, and *PRR7* was modified; *PRR9* and *PRR7* are now inhibited by *CCA1/LHY*. Also, a negative auto-regulation in *CCA1/LHY* was introduced as described in (Greenwood et al., 2022).

The resulting models consist of 11 ordinary differential equations (ODEs) to reproduce responses to light, rather than nine as in De Caluwé et al. 2016. Parameters values were taken from the original model. In model 2, where *PRR9* is repressed by *CCA1/LHY*, the mean of the original parameter values that go into *P*97 inhibition was used for parameterization; that is, the new parameter incorporated into the model is the mean between the Hill function parameters *K*_4_ and *K*_5_ (De Caluwé et al. 2016) for inhibition of *PRR9/PRR7* by *PRR5/TOC1* and *ELF4/LUX* in the original model, respectively. We name this parameter as *K*_11_. The added equations are,

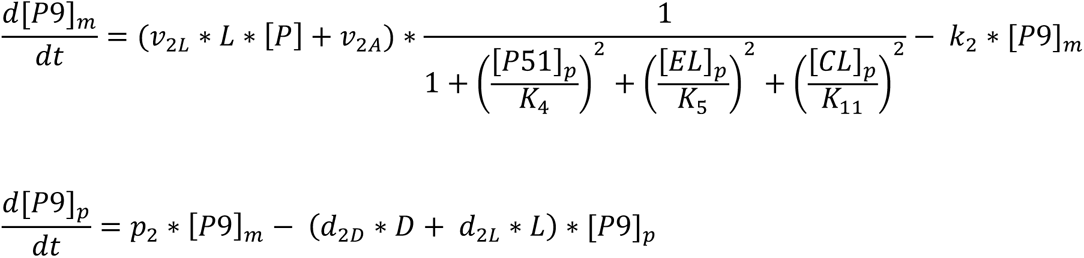

Simulations were carried out using experimental conditions, i.e., 8 days of entrainment at 12 hours light and 12 hours dark in a 24 h cycle. The clock then was released into constant light conditions. null mutants were simulated by setting the relevant rate constant of transcription to zero (*v_2a_* and *v_4_* in De Caluwé et al. 2016). All simulations were done in MATLAB.

### Luciferase circadian experiments

Seeds were surfaced sterilized before being sown on 1x MS plates with 3% w/v sucrose, 0.5 g/L MES and a pH of 5.7. Seeds were then stratified for three days. After three days, plates were moved to a neutral-day (ND) photoperiod chamber with a set temperature of 22°C and a light intensity of 85 μmol/m^-2^/s^-1^ for 5 days. On day 6, seedlings were transferred to a black 96 microwell plate containing the same media as described before. 15 μL of 5 mM luciferin was then added superficially to the media. Seedlings were then returned to the same entrainment chamber for 24-hours. After 24-hours of re-entrainment, seedlings were transferred to the TopCount before subjective dusk. All TopCount experiments were carried out under constant red and blue light. Data analysis was performed as described previously (Kolmos et al., 2009).

### Gene expression analysis

Comparative analysis of selected genes was performed by quantitative real-time PCR (qRT-PCR). Seedings were grown in a plant growth chamber (Percival Scientific, Model CU-36L5) at 22°C under 12:12 LD cycles (white light, 100 μmol/m^-2^/s^-1^) for 8 days and then transferred into constant light and temperature conditions. Samples were harvested and frozen at 3-h intervals. Total RNA was isolated using RNAiso Plus (TaKaRa, Cat. #9109) and reverse transcribed to produce cDNA using the RevertAid First Strand cDNA Synthesis Kit (Thermo Fisher Scientific, Cat. #K1622) following the manufacturer’s protocol. The qRT-PCR was performed by using the TB Green Premix Ex Taq (Tli RNase H Plus; TaKaRa, Cat. #RR420A) and CFX Connect Real-Time system (Bio-Rad). Primers sequences used for qRT-PCR are listed in table S1.

### Flowering time measurements

Flowering time experiments were carried out as described previously (Kolmos et al., 2011). In brief, seeds were surface sterilized and sown onto 1x MS plates with 0.25% w/v sucrose, 0.5 g/L MES and a pH of 5.7 and stratified for three days at 4°C. Plates were then moved to a neutral-day (ND) photoperiod chamber with a set temperature of 22°C and a light intensity of 85 μmol/m^-2^/s^-1^ for 14 days. After 14 days, seedlings were transferred to soil and moved to a short-day or long-day photoperiod. Under both photoperiods, the temperature was set to 22°C and light intensity of 85 μmol/m^-2^/s^-1^. Flowering was determined as the point at which the inflorescence was ∼1cm above the rosette. For each genotype, 15 to 20 plants were analyzed. All experiments were repeated twice with similar results observed each time. Statistical analysis was performed in Rstudio (v.2022.07.2) with the version 3.6.1 of R.

### Hypocotyl and root measurements

Seeds were surface sterilized and plated on solid *A. thaliana* solution (ATS) medium (Lincoln et al., 1990) with 1% w/v sucrose. Seeds were then stratified for three days before being transferred to a short-day chamber (8 hours of light, 16 hours of darkness) with a light intensity of 90 μmol/m^-2^/s^-1^. The temperature the seedlings were exposed to is described in text. Seedlings were allowed to grow for 8 days before scans of the plates were taken. The same scan was used to measure hypocotyl and root growth. Measurements were calculated using imageJ. A minimum of 42 seedlings were measured per genotype across two experimental repeats. Statistical analysis was performed in Rstudio (v.2022.07.2) with the version 3.6.1 of R.

## Supporting information

MATLAB code files

## Acknowledgements

This work was supported by funding from the UK Biotechnology and Biological Sciences Research Council (BSSRC): SJD: BB/N018540/1, DE/SJD: BB/V006665/1 and DE: BB/S506795/1. We also acknowledge BBSRC Whiterose DTP studentships (BB/M011151/1 and BB/T007222/1) to JR (ref 1792522) and EJW (ref 2444228). DE was also supported by the Royal Society (RGS\R2\212345). Deutsche Forschungsgemeinschaft (Qu 141/12-1) and the European Social Fund and the Federal State of Saxony-Anhalt (International Graduate School AGRIPOLY—Determinants of Plant Performance, grant no. ZS/2016/08/80644) supported MQ. PF was supported by the CONICYT PFCHA/DOCTORADO BECAS CHILE award (2013 – 72140562).

---

**Supplementary Figure 1.**
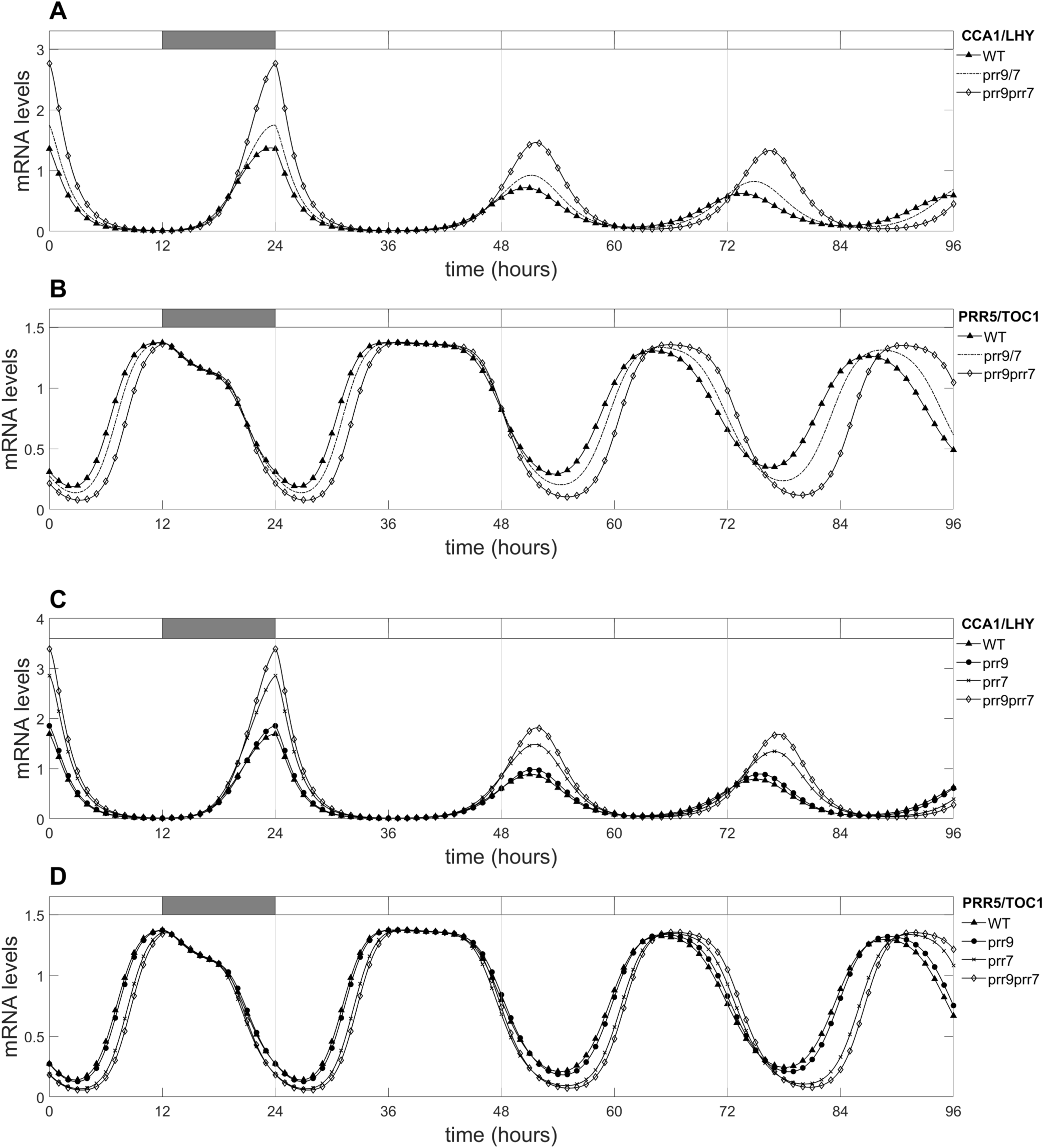
Simulating mutations in model 1 and model 2 replicates experimental data. To determine whether the modifications in (A) model 1 and (B) model 2 could replicate known circadian phenotypes, we simulated the *prr9*, *prr7* and *prr9/prr7* mutations by setting the respective gene’s transcription rate to zero. For model 1, the outputs for the *prr9* and *prr7* are combined into a single output (*prr9/7*) as model 1 keeps the same function for the respective single mutants.

**Supplementary Figure 2.**
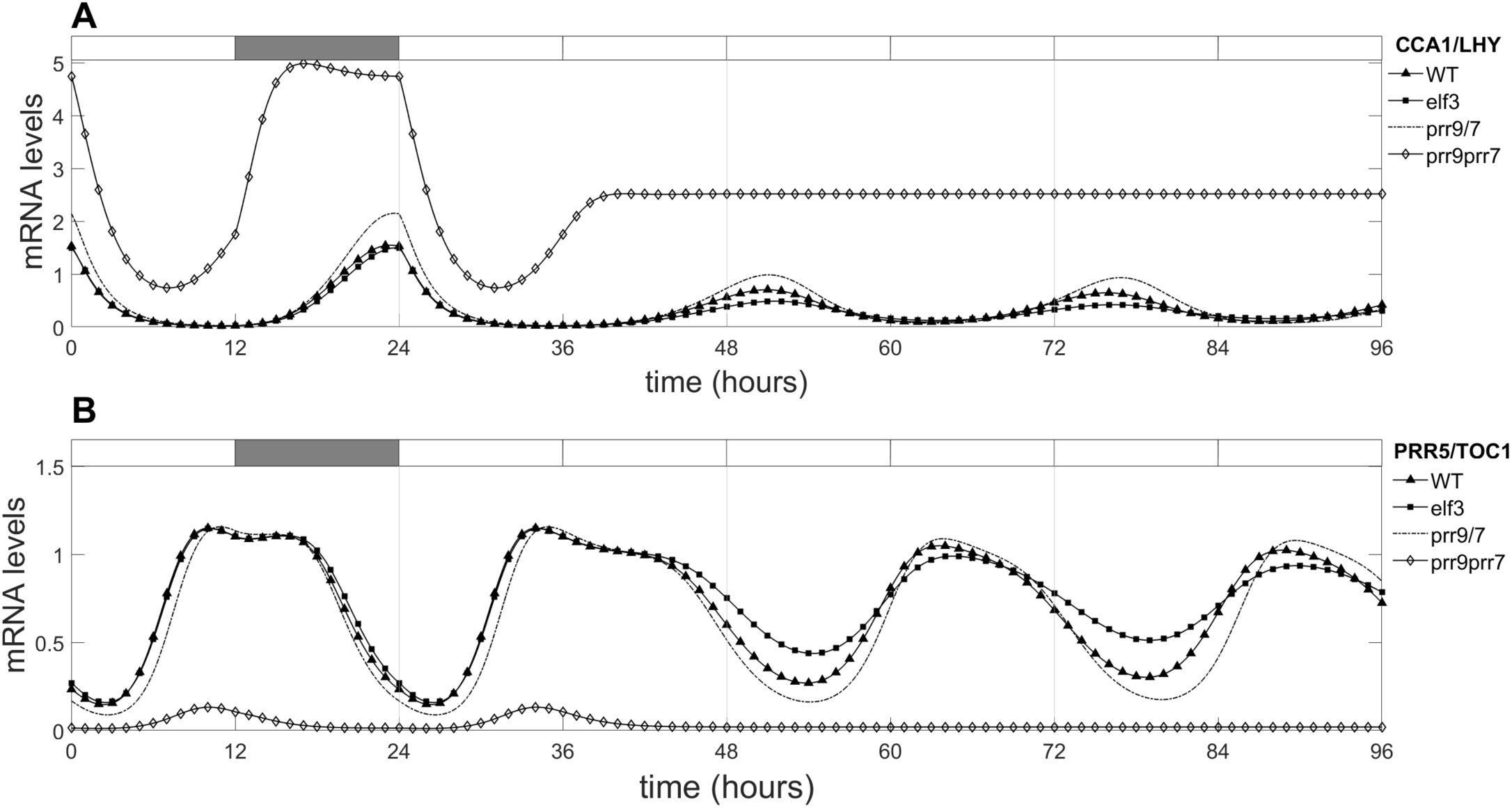
Model 3 does not replicate experimental data. In the third modification of the DC2016 model, we split the P97 component of the adaptation in Greenwood *et al*., 2022 into two separate components termed P9 and P7. The outputs of (A) *CL* and (B) *P51* in the WT and simulated *elf3/prr* mutant backgrounds are shown.

**Supplementary Figure 3.**
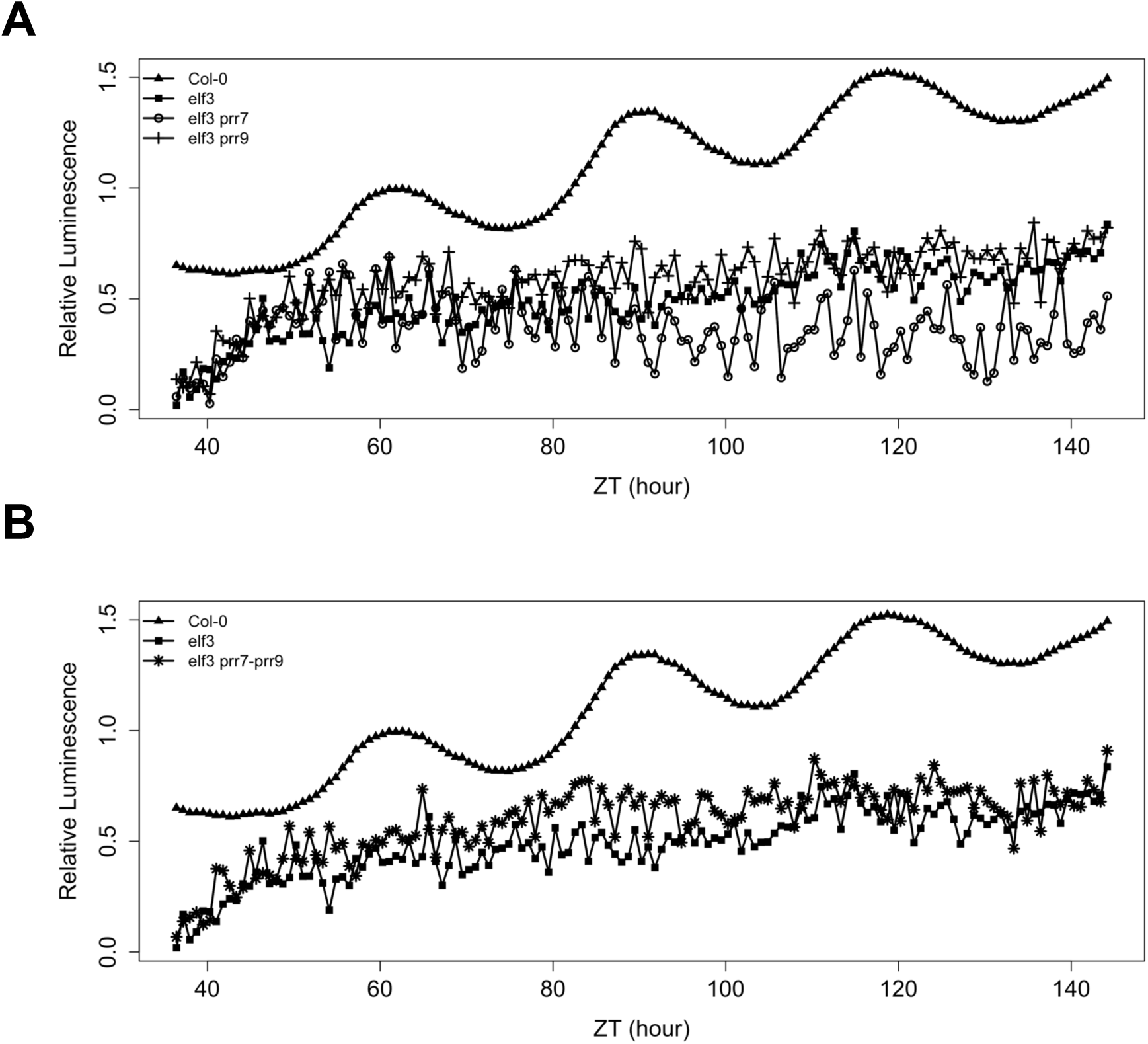
The *prr9* or *prr7* mutations does not rescue the arrhythmicity of the *CCA1::LUC* reporter in the *elf3* background. The arrhythmicity of the *CCA1::LUC* reporter in the *elf3-1* background is not restored by the (A) *prr9* or *prr7* single mutations. (B) The arrhythmicity of the *elf3 CCA1::LUC* was also not restored by simultaneous mutations in *prr9 and prr7*. Seedlings were entrained under neutral-day (12/12) cycles before being released into constant light and temperature. The first day under constant conditions is not shown.

**Supplementary Figure 4.**
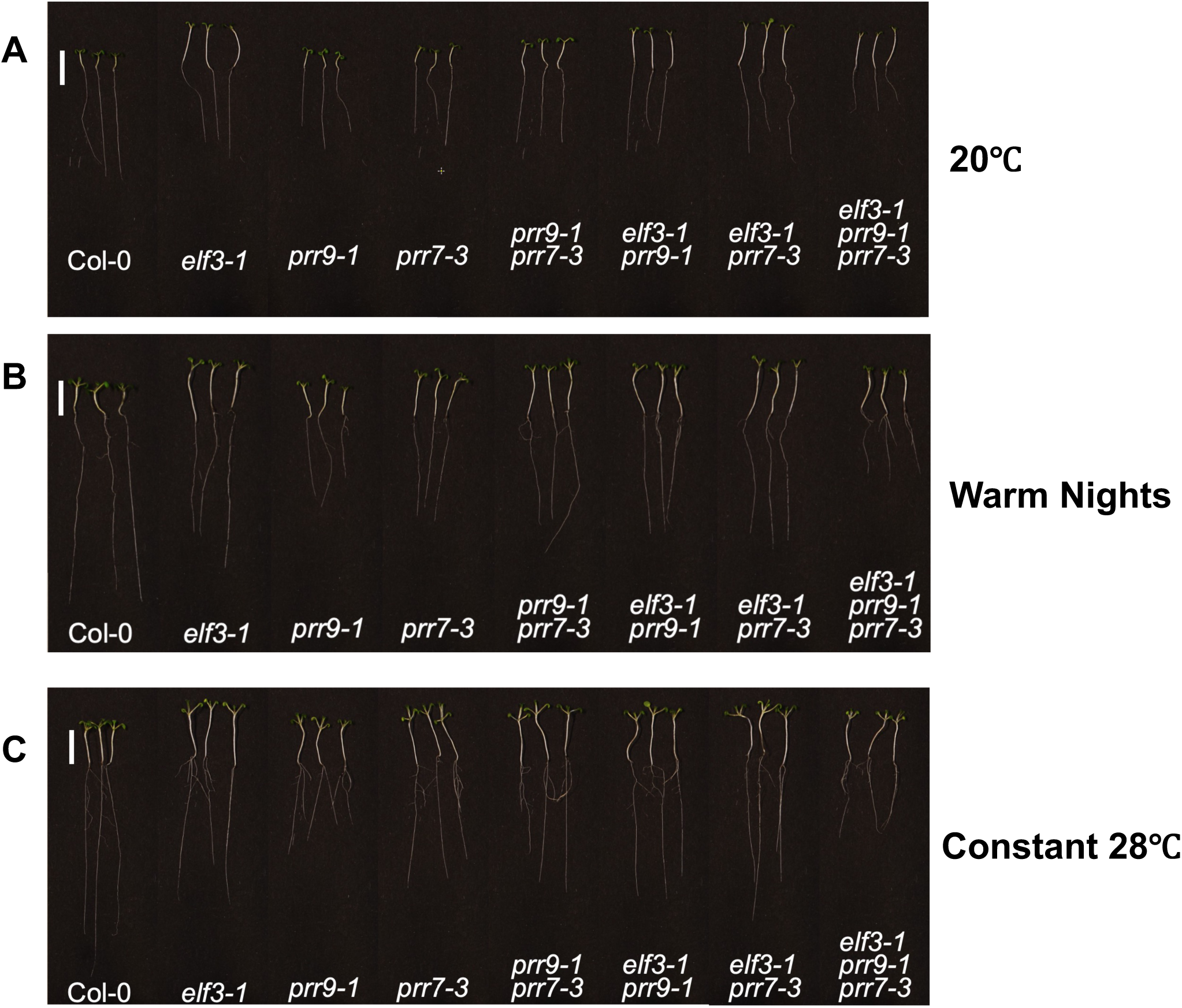
Representative images of the *elf3* and *prr* lines. Images of seedlings grown under short-day photoperiods with (A) 20°C, (B) warm 28°C nights only and (C) constant warm (28°C) temperatures. Scale bars are 10 mm.

**Supplementary table 1.**
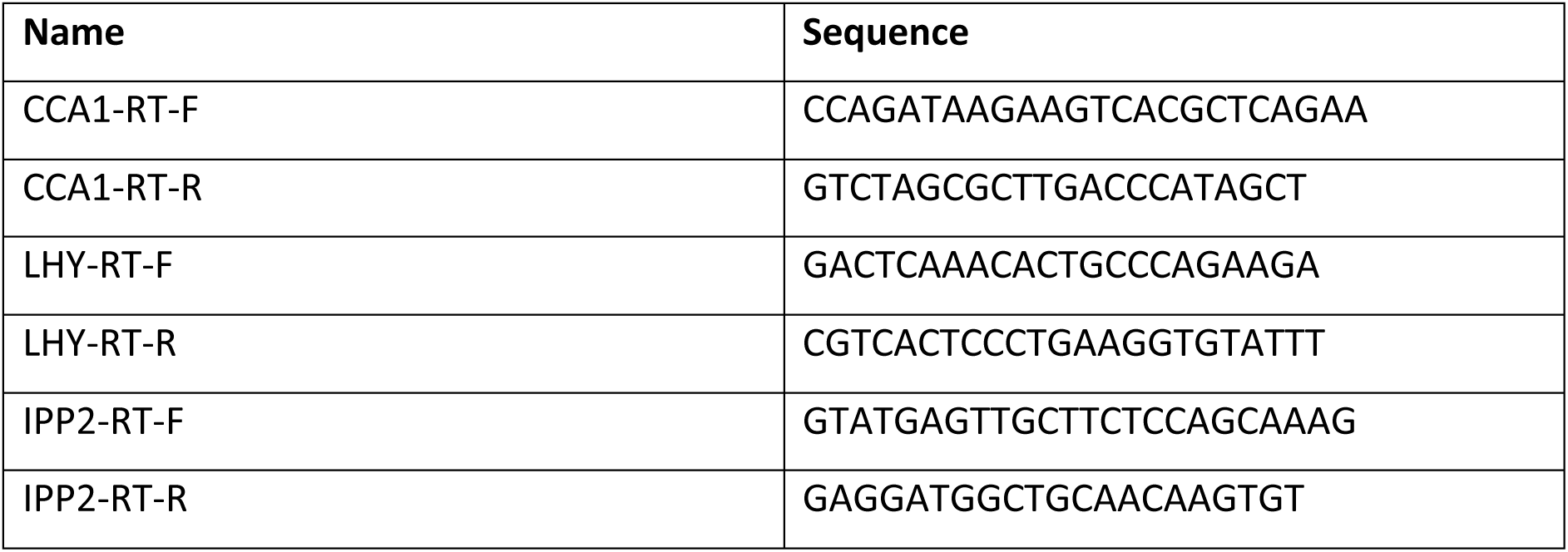
qPCR primers used in this work.

