## Supplementary material for "Complex epistatic interactions between ELF3, PRR9, and PRR7 regulates the circadian clock and plant physiology": MATLAB code files: DC2016model3.m

```
function F = DC2016model3(t,y)
% the modified DC2016 model in Greenwood et al. 2012 (single-cell model)
% PRR9/PRR7 is split
% y will be a vector holding the mRNA and protein levels of the main clock genes
% grouped into: CL, P9, P7, P51, EL and the dark-accumulating protein P
% t is time
% y(1)  CL  mRNA
% y(2)  CL  protein
% y(3)  P9  mRNA
% y(4)  P9  protein
% y(5)  P7  mRNA
% y(6)  P7  protein
% y(7)  P51 mRNA
% y(8)  P51 protein
% y(9)  EL  mRNA
% y(10) EL  protein
% y(11) P   protein

global  L D v1 v1L v2A v2L v3 v4 k1L k1D k2 k3 k4 p1 p1L p2 p3 p4 d1 d2D d2L d3D d3L d4D d4L K1 K2 K4 K5 K6 K7 K8 K9 K10 K0 K5b v2AA;
        % parameters for the core clock model
        
dCLmdt = (v1 + v1L * L * y(11)) * (1/(1+ (y(2)/K0)^2 +(y(4)/K1)^2 +(y(6)/K1)^2+ (y(8)/K2)^2)) - (k1L * L + k1D * D) * y(1);
dCLpdt = (p1 + p1L * L) * y(1) - d1 * y(2);
dP9mdt = (v2L * L * y(11) + v2AA) * (1 / (1 + ((y(8)/K4)^2) + ((y(10)/K5)^2) + ((y(2)/K5b)^2) )) - k2 * y(3);
dP9pdt =  p2 * y(3) - (d2D * D + d2L * L) * y(4);
dP7mdt = (v2L * L * y(11) + v2A) * (1 / (1 + ((y(8)/K4)^2) + ((y(10)/K5)^2) + ((y(2)/K5b)^2) )) - k2 * y(5);
dP7pdt =  p2 * y(5) - (d2D * D + d2L * L) * y(6);
dP51mdt =  v3 * ( 1 / (1 + (y(2)/K6)^2 + (y(8)/K7)^2)) - k3 * y(7);
dP51pdt =  p3 * y(7) - (d3D * D + d3L * L) * y(8);
dELmdt  =  L * v4 * (1/(1 + (y(2)/K8)^2 + (y(8)/K9)^2 + (y(10)/K10)^2 )) - k4 * y(9);
dELpdt =  p4 * y(9) - (d4D * D + d4L * L) * y(10);
dPdt   =  0.3 * (1 - y(11)) * D - (y(11) * L);

         F = [dCLmdt ;dCLpdt ;dP9mdt ;dP9pdt ;dP7mdt ;dP7pdt; dP51mdt ;dP51pdt ;dELmdt ;dELpdt ;dPdt];
end


        
        
```
