## Supplementary material for "Complex epistatic interactions between ELF3, PRR9, and PRR7 regulates the circadian clock and plant physiology": MATLAB code files: DC2016model_modifications.m

```
% CLOCK MODEL for light entrainment (De Caluwé et al. 2016)

% De original DC2016 model is modified by splitting P97 component into P9 (PRR9) and P7 (PRR7) components
% DC2016model1 maintains the roles of the model components as originally set in De Caluwé et al. 2016
% DC2016model2 changes the original promoter role of CL on P9 to be inhibitor
% DC2016model3 is the DC2016 adaptation in Greenwood et al. 2022
% Null mutants are simulated by setting the relevant synthesis rates to zero

close all;
clear all;


global  L D v1 v1L v2A v2B v2L v3 v4 k1L k1D k2 k3 k4 p1 p1L p2 p3 p4 d1 d2D d2L d3D d3L d4D d4L K1 K2 K3 K4 K5 K6 K7 K8 K9 K10 v2AA;


% Parameters values -----------------------------------------------
run parameters_model1_and_model2
% run parameters_model3

% v2AA   =  0; % P9 synthesis ------------------------ NULL MUTANTS
% v2A	 = 	0; % P7 synthesis ------------------------ NULL MUTANTS
% v4	 = 	0; % EL synthesis ------------------------ NULL MUTANTS


% entrainment of the clock for a 24 h light/dark cycle

lighttime=12; % hours in light
darktime=24-lighttime; % hours in dark
totaltime=8*24; % total time in entrained conditions
elapsed=0;
ystart=rand(1,11)*0.1; % random initial values

while elapsed<totaltime
    % in light
    L=1;
    D=0;
    [t,y]=ode23s(@DC2016model1,[elapsed:0.01:elapsed+lighttime],ystart);
    elapsed=elapsed+lighttime;
    ystart= y(end,:);
    subplot(2,1,1)
    plot(t,y(:,1), 'linewidth',1.5, 'Color', 'k');
    hold on
    subplot(2,1,2)
    plot(t,y(:,7), 'linewidth',1.5, 'Color', 'k');
    hold on
    
    % in dark
    L=0;
    D=1;
    [t,y]=ode23s(@DC2016model1,[elapsed:0.01:elapsed+darktime],ystart);
    elapsed=elapsed+darktime;
    ystart= y(end,:);
    subplot(2,1,1)
    plot(t,y(:,1), 'linewidth',1.5, 'Color', 'k');
    subplot(2,1,2)
    plot(t,y(:,7), 'linewidth',1.5, 'Color', 'k');
    
          
end


% constant light conditions
     L = 1;
     D = 0;
     ystart= y(end,:);
     [t,y]=ode23s(@DC2016model1,[elapsed:0.01:elapsed+3*24],ystart);

% plotting the last 24 hours under entrained conditions (12L:12D) followed by 72 hours in constant light               
subplot(2,1,1)
plot(t,y(:,1),'linewidth',1.5,'Color', 'k');
set(gca,'FontSize',16);
ylabel('CCA1/LHY mRNA','fontSize',22)
xlabel('time (hours)','fontSize',26)
set(gca, 'XTick', 0+7*24:12:11*24);
xlim([0+7*24 11*24])
my_labels=[{'0','12','24','36','48','60','72','84','96','108','120','132','144','156','168','180','192','204','216','228','240','252','264'}];
set(gca,'XTickLabel',my_labels);

subplot(2,1,2)
plot(t,y(:,7),'linewidth',1.5,'Color', 'k');
set(gca,'FontSize',16);
ylabel('PRR5/TOC1 mRNA','fontSize',22)
xlabel('time (hours)','fontSize',26)
set(gca, 'XTick', 0+7*24:12:11*24);
xlim([0+7*24 11*24])
my_labels=[{'0','12','24','36','48','60','72','84','96','108','120','132','144','156','168','180','192','204','216','228','240','252','264'}];
set(gca,'XTickLabel',my_labels);
```
