## Supplementary material for "Complex epistatic interactions between ELF3, PRR9, and PRR7 regulates the circadian clock and plant physiology": MATLAB code files: parameters_model1_and_model2.m

```
% it sets the parameters values for model 1 and model 2

global  L D v1 v1L v2A v2B v2L v3 v4 k1L k1D k2 k3 k4 p1 p1L p2 p3 p4 d1 d2D d2L d3D d3L d4D d4L K1 K2 K3 K4 K5 K6 K7 K8 K9 K10 v2AA;


% Parameters values in De Caluwé et al. 2016
 
v1	 = 	4.60; % CL synthesis
v1L	 = 	3.00; % CL light-induced systhesis
v2AA = 	1.30; % P9 synthesis (PRR9)
v2A	 = 	1.30; % P7 synthesis (PRR7)
v2B	 = 	1.50; % P9 and P7 CL-induced synthesis
v2L	 = 	5.00; % P9 and P7 light-induced synthesis
v3	 = 	1.00; % P51 synthesis
v4	 = 	1.50; % EL synthesis
k1L	 = 	0.53; % CL mRNA degradation (light) 
k1D	 = 	0.21; % CL mRNA degradation (dark)
k2	 = 	0.35; % P9 and P7 mRNA degradation
k3	 = 	0.56; % P51 mRNA degradation
k4	 = 	0.57; % EL mRNA degradation
p1	 = 	0.76; % CL translation
p1L	 = 	0.42; % CL light light-induced translation
p2	 = 	1.00; % P9 and P7 translation
p3	 = 	0.64; % P51 translation
p4	 = 	1.00; % EL translation
d1	 = 	0.68; % CL degradation
d2D	 = 	0.50; % P9 and P7 degradation (dark)
d2L	 = 	0.30; % P9 and P7 degradation (light)
d3D	 = 	0.48; % P51 degradation (dark)
d3L	 = 	0.78; % P51 degradation (light)
d4D	 = 	1.20; % EL degradation (dark)
d4L	 = 	0.38; % EL degradation (light)
K1	 = 	0.16; % Inhibition: CL by P9 and P7
K2	 = 	1.20; % Inhibition: CL by P51
K3	 = 	0.24; % Activation: P9 and P7 by CL
K4	 = 	0.23; % Inhibition: P9 and P7 by P51
K5	 = 	0.30; % Inhibition: P9 and P7 by EL
K6	 = 	0.46; % Inhibition: P51 by CL
K7	 = 	2.00; % Inhibition: P51 by itself
K8	 = 	0.36; % Inhibition: EL by CL
K9	 = 	1.90; % Inhibition: EL by P51
K10	 = 	1.90; % Inhibition: EL by EL
```
