## Supplementary material for "Complex epistatic interactions between ELF3, PRR9, and PRR7 regulates the circadian clock and plant physiology": MATLAB code files: parameters_model3.m

```
% it sets the parameters values for model 3

global  L D v1 v1L v2A v2L v3 v4 k1L k1D k2 k3 k4 p1 p1L p2 p3 p4 d1 d2D d2L d3D d3L d4D d4L K1 K2 K4 K5 K6 K7 K8 K9 K10 K0 K5b v2AA;


% Parameters values in De Caluwé et al. 2016
 
v1	 = 	4.58; % CL synthesis                    
v1L	 = 	3.00; % CL light-induced systhesis
v2AA =  1.27; % P97 synthesis (PRR9)            
v2A	 = 	1.27; % P97 synthesis (PRR7)            
v2L	 = 	5.00; % P97 light-induced synthesis
v3	 = 	1.00; % P51 synthesis
v4	 = 	1.47; % EL synthesis                    
k1L	 = 	0.53; % CL mRNA degradation (light) 
k1D	 = 	0.21; % CL mRNA degradation (dark)
k2	 = 	0.35; % P97 mRNA degradation
k3	 = 	0.56; % P51 mRNA degradation
k4	 = 	0.57; % EL mRNA degradation
p1	 = 	0.76; % CL translation
p1L	 = 	0.42; % CL light light-induced translation
p2	 = 	1.01; % P97 translation                  
p3	 = 	0.64; % P51 translation
p4	 = 	1.01; % EL translation                   
d1	 = 	0.68; % CL degradation
d2D	 = 	0.50; % P97 degradation (dark)
d2L	 = 	0.29; % P97 degradation (light)          
d3D	 = 	0.48; % P51 degradation (dark)
d3L	 = 	0.38; % P51 degradation (light) --(changed by Greenwood et al. 2022)
d4D	 = 	1.21; % EL degradation (dark)           
d4L	 = 	0.38; % EL degradation (light)
K0   =  2.80; % Inhibition: CL by CL --(introduced by Greenwood et al. 2022)
K1	 = 	0.16; % Inhibition: CL by P97
K2	 = 	1.18; % Inhibition: CL by P51           
K4	 = 	0.28; % Inhibition: P97 by P51 --(changed by Greenwood et al. 2022)
K5	 = 	0.57; % Inhibition: P97 by EL --(changed by Greenwood et al. 2022)
K5b  =  1.73; % Inhibition: P97 by CL --(introduced by Greenwood et al. 2022)
K6	 = 	0.46; % Inhibition: P51 by CL
K7	 = 	2.00; % Inhibition: P51 by itself
K8	 = 	0.36; % Inhibition: EL by CL
K9	 = 	1.90; % Inhibition: EL by P51
K10	 = 	1.90; % Inhibition: EL by EL
```
