## Supplementary material for "Complex epistatic interactions between ELF3, PRR9, and PRR7 regulates the circadian clock and plant physiology": MATLAB code files: Readme.docx

We implemented three modifications of the De Caluwé et al. (2016) model (model 1, model 2, and model 3). The equations of these models are in the files DC2016model1.m, DC2016model2.m, and DC2016model3.m. The file parameters_mode1_and_model2.m defines the parameter values for model 1 and model 2; and the file parameters_model3 defines the parameter values for model 3. The script to execute the models is in the file DC2016model_modifications.m.

**Wild type simulations**

In DC2016model_modifications.m (lines 38, 51, and 67), define the model to be executed by setting the relevant function (e.g. for model 1, the function DC2016model1.m). Set the parameter values for the relevant model in lines 18 or 19.

**Simulating clock mutants**

In DC2016model_modifications.m, after setting the parameter values for a model (in lines 18 or 19), specify the mutant to be produced by changing the relevant parameter value in lines 21, 22, and/or 23.
